# Revisiting the diversity of secondary endosymbionts in the major pest oat aphid, *Rhopalosiphum padi*

**DOI:** 10.64898/2026.05.19.726398

**Authors:** Qiong Yang, Bin Zhu, Wenjuan Yu, Zhengyang Zhao, Alex Gill, Jasmeen Kaur, Nadieh de Jonge, Jun-Bo Luan, Torsten N. Kristensen, Pei Liang, Ary A. Hoffmann

**Affiliations:** Pest and Environmental Adaptation Research Group, School of BioSciences, The University of Melbourne, Parkville, VIC 3052, Australia; Department of Entomology, College of Plant Protection, China Agricultural University, China; Ministry of Agriculture Key Laboratory of Integrated Management of Pests on Crops in Southwest China, Institute of Plant Protection, Sichuan Academy of Agricultural Sciences, Chengdu, China; Liaoning Key Laboratory of Economic and Applied Entomology, College of Plant Protection, Shenyang Agricultural University, Shenyang, China; Section for Functional Ecology and Genomics, Department of Chemistry and Bioscience, Aalborg University, Aalborg 9220, Denmark

## Abstract

There is disagreement on whether secondary endosymbionts are found in the major cereal pest aphid, *Rhopalosiphum padi*. Some papers report a diversity of secondary bacterial endosymbionts while others have failed to find evidence of these bacteria in this species. Here we revisit this issue by summarizing the relevant literature and through additional sampling of the species in Australia, China and Denmark using a combination of molecular approaches. We find a general absence of secondary endosymbionts beyond the obligate endosymbiont *Hamiltonella defensa* in *R. padi*. While the inconsistency in survey results may reflect rapid changes in endosymbiont turnover in populations and/or the impact of ecological factors such as host plant type on endosymbiont diversity, we are concerned that technical issues may be at least partly responsible for inconsistencies in the literature. This leads us to emphasize the importance of multiple sources of evidence required to establish and characterize endosymbiont infections, including PCR and qPCR assays, DNA Sanger sequencing and 16SrRNA gene metabarcoding. We note that several major aphid pests show a low incidence of secondary endosymbionts which raises issues about the importance of these endosymbionts in aphids that constitute pests, even though endosymbionts can in some cases increase host fitness and therefore pest impact.

## Introduction

Aphids can contain a range of bacteria, several of which occur in the body of the aphids outside gut tissue and are passed on through generations vertically (Moran et al., 2008). There is ongoing interest in understanding the role of these endosymbiotic bacteria on their aphid hosts, both from the perspective of aphid control and from understanding host-bacterial interactions on aphid fitness effects. Endosymbionts can increase host fitness (and potentially therefore pest status) (Zytynska et al., 2021) and the ability of pests to be affected by parasitoids. Endosymbionts are often seen as providing “protection” against parasitoids although traits related to protection can be decreased (Luo et al., 2020), have no effect (Hansen et al., 2012; Lenhart & White, 2017), or increased (Oliver et al., 2003) depending on the bacterial-host combination. Two endosymbionts, *Hamiltonella defensa* (Ayoubi et al., 2025; Oliver et al., 2003) and *Regiella insecticola* (Vorburger et al., 2010), are considered particularly important for providing protective functions which could diminish biological control by parasitoids and favour pest outbreaks (Vorburger, 2014). However other strains of related or different endosymbionts may benefit pest control such as by decreasing plant virus transmission (Yu et al., 2025) or increasing pesticide susceptibility (Skaljac et al., 2018). For these reasons, it is important to accurately establish the distribution of bacterial endosymbionts in major pest aphids and their potential phenotypic effects on hosts.

The oat aphid, *Rhopalosiphum padi*, is regarded as a major pest aphid in many parts of the world. The species is responsible for direct damage of wheat and other cereal crops and moreover can transmit plant viruses such as strains of Barley Yellow Dwarf Virus which can be devastating (Walls III et al., 2019; Zhao & Zhou, 2025). Our initial interest in the endosymbionts of *R. padi* stemmed from the fact that in initial surveys in Australia we failed to find any evidence of secondary endosymbionts in this species (Yang et al., 2023a). This finding stood in stark contrast to findings reported from China and Europe where a number of secondary endosymbionts had been reported at a high apparent incidence, including *Hamiltonella, Regiella*, and *Arsenophonus* (Guo et al., 2019; Majeed et al., 2022).

Such differences in survey results may have several explanations. One possibility is that endosymbiont incidence can vary among populations of clonal lineages (Leybourne, 2025), which is particularly pertinent to Australia where aphids are regarded as invasive (Yang et al., 2023a); the invasive process is expected to decrease clonal (and any associated endosymbiont) diversity. Another possibility is that endosymbiont incidence varies seasonally or by host plant, both of which have been documented for various endosymbionts (Csorba et al., 2024; Vorburger, 2014). However, it is also possible that other issues are involved including incorrect detections of endosymbionts in some studies, particularly if controls are inadequate, if there is a risk of contamination, and/or if inappropriate techniques are used to detect the endosymbionts. Such issues have for instance re-occurred in the detection of *Wolbachia* endosymbionts in *Aedes aegypti* mosquitoes (Ross et al., 2020).

These concerns have led us to reinvestigate the presence of bacterial endosymbionts in *R. padi*, which is relevant for exploring the utility of endosymbiont transinfections in helping to control this pest and its ability to transmit plant viral diseases (Yu et al., 2025). Any efforts based on transinfections may be restricted in their effectiveness if there are already diverse endosymbionts present in the pest species that vary with geographical and ecological factors. To develop an understanding of the distribution of endosymbionts in *R. padi*, we undertake (1) a literature review of relevant studies and the approaches used for detection, (2) additional surveys of the species in Australia from different host plants, (3) new surveys of the distribution of endosymbionts in samples from China and Denmark, and (4) a comparison of approaches used for detecting the endosymbionts. While it is difficult to make firm conclusions about the reasons for inconsistencies in detection of bacterial endosymbionts in this species, we argue that a higher level of technical rigour could be useful in validating detections.

## Methods

### Literature review

We searched for papers that mentioned “*Rhopalosiphum padi*” and “endosymbiont” or any of the endosymbionts recorded in *R. padi* (below) based on searches in Google Scholar and Web of Science (completed 24 December 2025). We then recorded the geographic area covered and the nature of the molecular approach(es) used for detection. In some cases, single pooled samples were available from the same field population while in others multiple samples or individuals were individually tested from the same population (Table 1). Plant type and controls were also recorded for samples where provided.

**Table 1.**
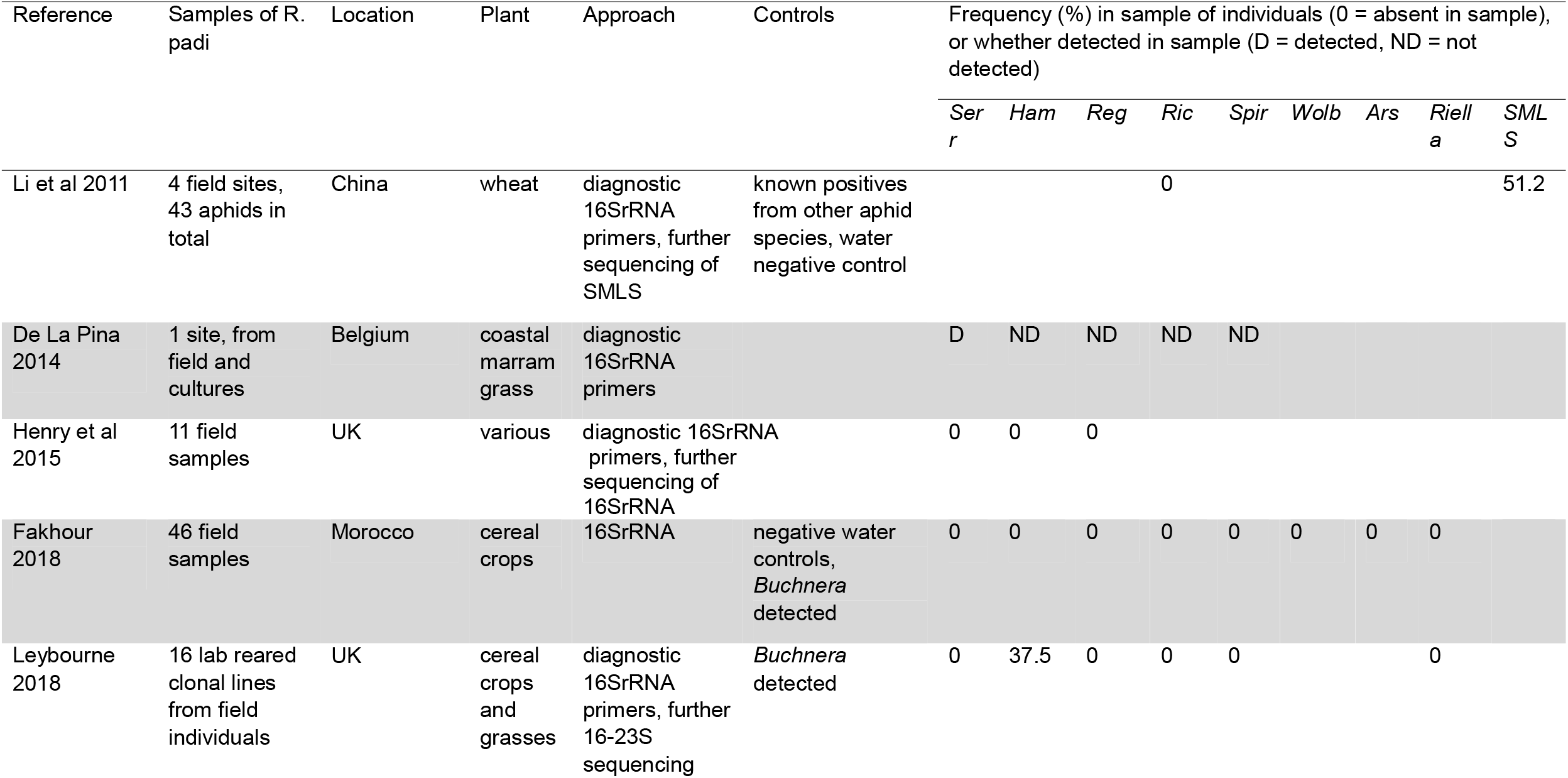
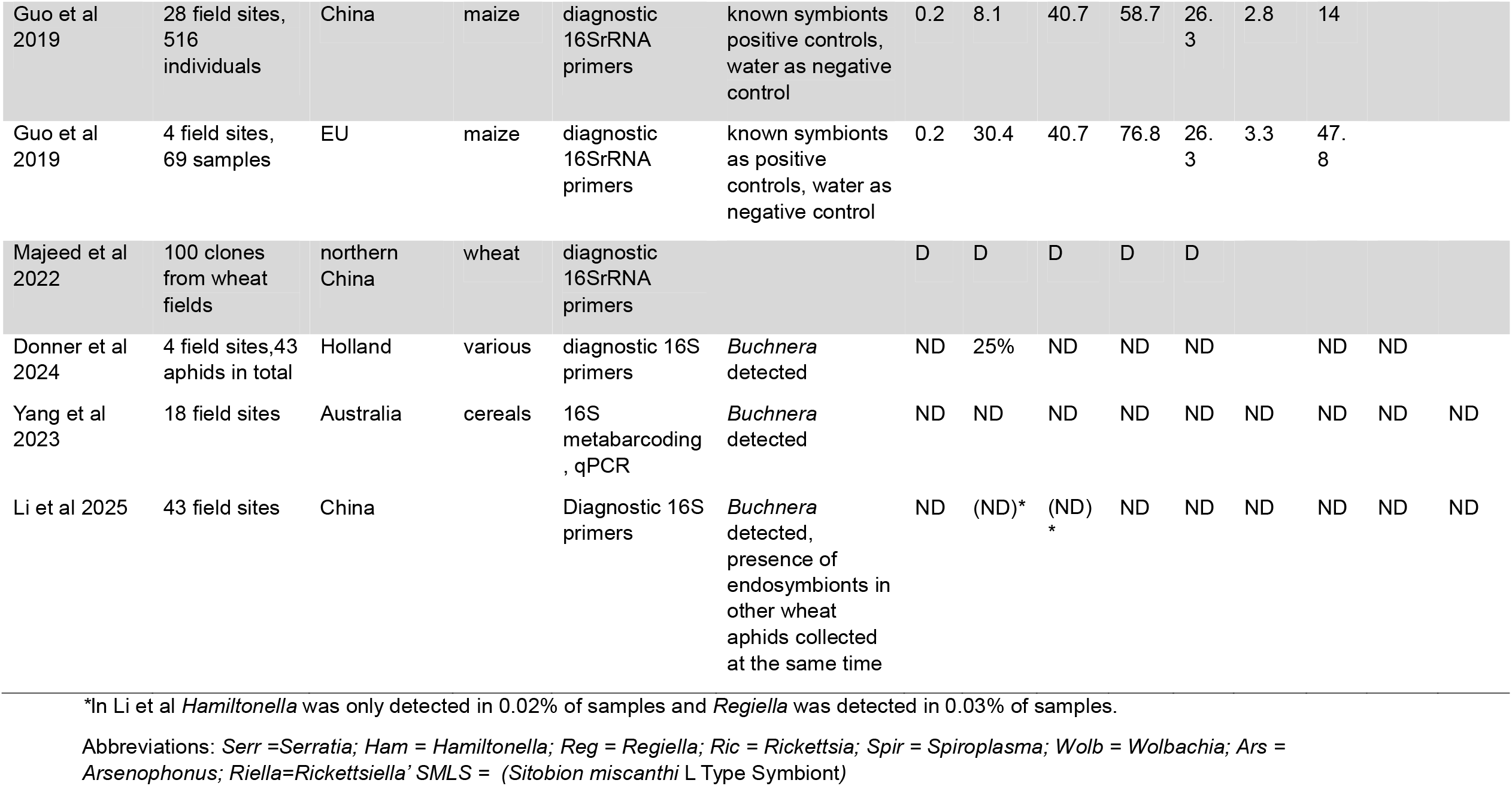
Results of literature survey on distribution of R. padi detections of secondary endosymbionts.

### New aphid samples

Forty-eight *R. padi* samples were collected by direct searching for aphids from a variety of host plants from around Australia between 2019 and 2026, while several historical samples, which were collected from the field and had been placed in culture, were also included (Table 2). Ten individuals from each sample were stored in 100% ethanol and frozen at −20°C for later molecular analysis. The samples in culture were maintained as asexual lines established from a single female in the laboratory on excised wheat seedlings (var. Trojan)14 days after sowing (Zadoks GS12), that were inserted into cups containing a water reservoir. All aphid cultures were kept in a controlled temperature (CT) room at 19 ± 1 °C with a 16:8 h light:dark photoperiod. Fifty-nine samples were collected from China from wheat, barley or corn between 2023 and 2025 (Table 3). An additional four samples from Denmark were included in this study (Table 4). Maps displaying the location of all *R. padi* samples used are provided in Figures 1-3. For comparison with another aphid found in the same crop type as *R. padi*, some collections of *Sitobion avenae* aphids were also made from China and a single collection of *S. fragariae* from Australia (Table 5).

**Table 2.**
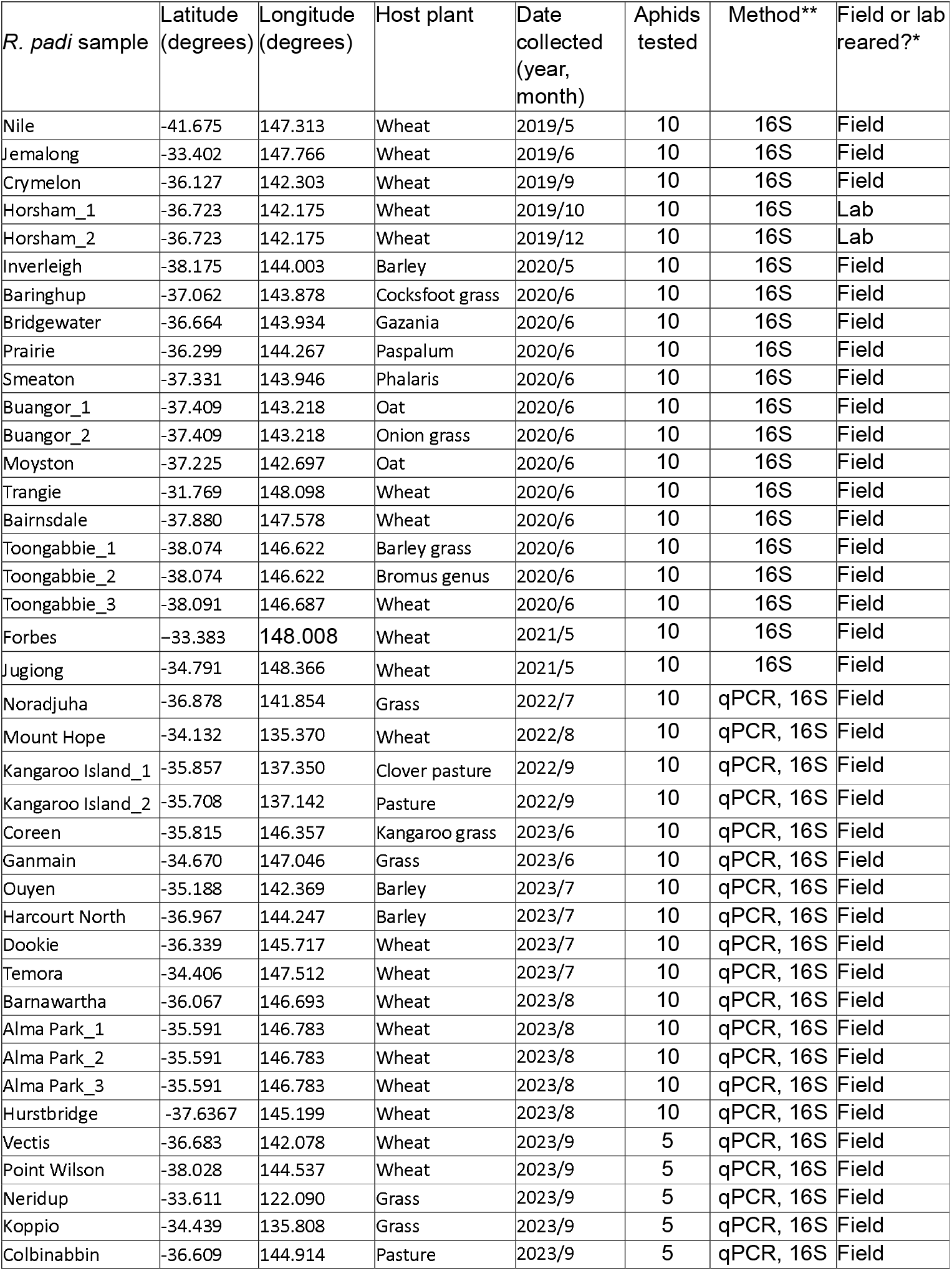
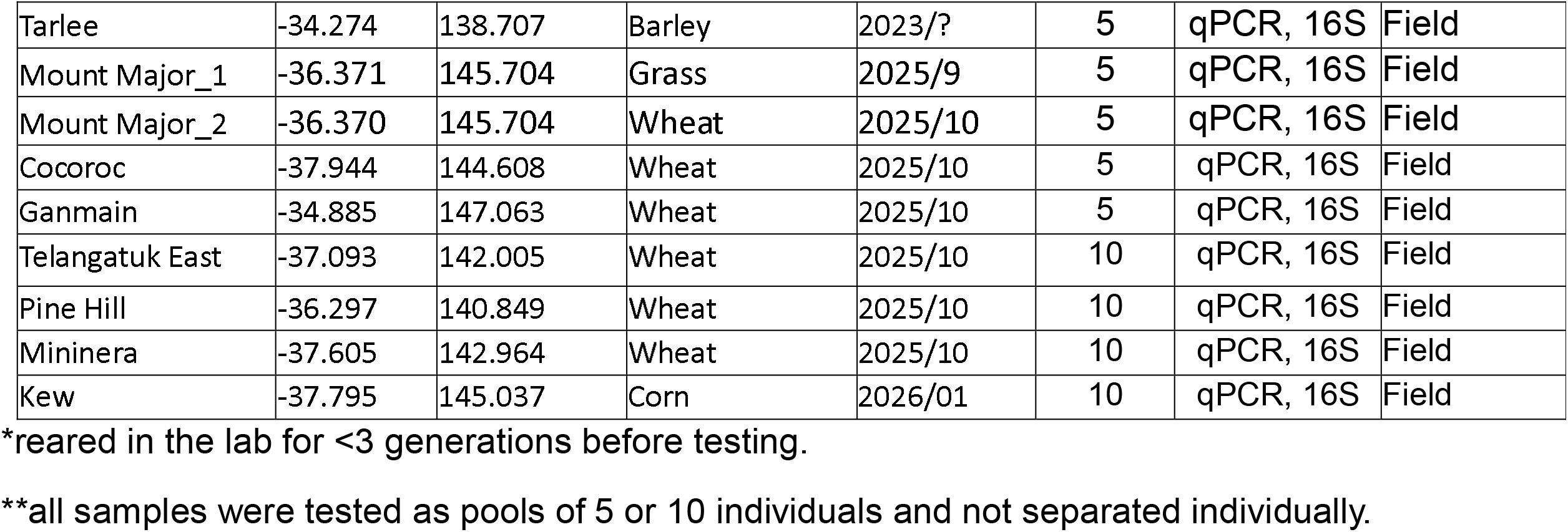
Information on field samples of *R. padi* collected from Australia. Samples tested were taken from the field or after <3 generations of lab rearing. No secondary endosymbionts were found. For mapped locations see Figure 1.

**Table 3.** Information on field samples of *R. padi* collected from China as mapped in Figure 2. Samples tested were taken directly from the field or after being reared in the laboratory for 3 generations. No secondary endosymbionts were found.

| *R. padi* sample | Latitude (degrees) | Longitude (degrees) | Host plant | Date Collected (year, month) | Aphids tested | Method | Field or lab reared? |
| --- | --- | --- | --- | --- | --- | --- | --- |
| Baoding | 38.850000 | 115.483333 | Corn | 2025/9 | 10 | qPCR, 16S | Field |
| Baoshan | 25.114861 | 99.160750 | Barley | 2024/5 | 10 | qPCR, 16S | Lab |
| Bengbu | 32.910000 | 117.240000 | Barley | 2025/4 | 10 | PCR, 16S | Field |
| Dali | 25.670000 | 100.300000 | Barley | 2025/3 | 10 | PCR, 16S | Field |
| Dayi | 30.470836 | 103.572284 | Wheat | 2024/3 | 10 | 16S | Field |
| Deyang | 30.977062 | 104.282492 | Wheat | 2024/3 | 10 | 16S | Field |
| Ezhou | 30.010000 | 114.910000 | Barley | 2025/5 | 10 | PCR, 16S | Field |
| Gushi | 32.171167 | 115.648889 | Wheat | 2024/5 | 10 | qPCR, 16S | Lab |
| Haibei | 36.958194 | 100.901306 | Barley | 2024/7 | 10 | qPCR, 16S | Lab |
| Haidian_4 | 39.950000 | 116.283333 | Corn | 2025/9 | 10 | qPCR, 16S | Field |
| Haidian_1 | 39.906222 | 116.391278 | Wheat | 2024/5 | 10 | qPCR, 16S | Lab |
| Haidian_2 | 39.950000 | 116.290000 | Wheat | 2025/5 | 10 | PCR, 16S | Field |
| Haidian_3 | 40.030000 | 116.290000 | Wheat | 2025/5 | 10 | PCR, 16S | Field |
| Handan_1 | 36.550000 | 114.410000 | Wheat | 2023/5 | 10 | 16S | Lab |
| Handan_2 | 36.550000 | 114.410000 | Wheat | 2025/5 | 10 | PCR, 16S | Field |
| Hangzhou_1 | 30.248972 | 120.205222 | Barley | 2024/4 | 10 | qPCR, 16S | Lab |
| Hangzhou_2 | 30.248972 | 120.205222 | wheat | 2025/5 | 10 | PCR, 16S | Field |
| Hefei_1 | 31.853111 | 117.255028 | Wheat | 2023/5 | 10 | 16S | Lab |
| Hefei_2 | 31.853111 | 117.255028 | Wheat | 2024/4 | 10 | qPCR, 16S | Lab |
| Hefei_3 | 31.820000 | 117.220000 | Barley | 2025/5 | 10 | PCR, 16S | Field |
| Hohhot | 40.833806 | 111.673083 | Barley | 2024/7 | 10 | qPCR, 16S | Lab |
| Huaian | 33.750000 | 118.840000 | Wheat | 2023/5 | 10 | 16S | Field |
| Huainan | 32.586694 | 117.013000 | Wheat | 2024/4 | 10 | qPCR, 16S | Lab |
| Jiangyou_1 | 31.892624 | 104.788344 | Wheat | 2024/4 | 10 | PCR, 16S | Field |
| Jiangyou_2 | 31.872200 | 104.802500 | Wheat | 2024/4 | 10 | 16S | Field |
| Jiaxing | 30.730000 | 120.780000 | Wheat | 2023/5 | 10 | 16S | Lab |
| Jiaxing_1 | 30.747444 | 120.751083 | Wheat | 2024/5 | 10 | qPCR, 16S | Lab |
| Jiaxing_2 | 30.747444 | 120.751083 | Wheat | 2025/3 | 10 | PCR, 16S | Field |
| Jingmen | 31.110000 | 112.260000 | Wheat | 2023/5 | 10 | 16S | Lab |
| Jining | 35.540000 | 116.670000 | Wheat | 2023/5 | 10 | 16S | Lab |
| Langfang | 39.420000 | 116.380000 | Wheat | 2025/5 | 10 | PCR, 16S | Field |
| Lanping | 26.502400 | 99.222682 | Wheat | 2024/3 | 10 | 16S | Field |
| Luoyang | 34.619667 | 112.447694 | Wheat | 2024/5 | 10 | qPCR, 16S | Lab |
| Ningnan | 27.073623 | 102.747862 | Wheat | 2024/4 | 10 | 16S | Field |
| Pixian_1 | 30.812155 | 103.849430 | Wheat | 2024/4 | 10 | PCR, 16S | Field |
| Pixian_2 | 30.810000 | 103.850000 | Wheat | 2024/2 | 10 | PCR, 16S | Lab |
| Pixian_3 | 30.670000 | 104.060000 | Wheat | 2025/2 | 10 | qPCR, 16S | Field |
| Pixian_4 | 30.670000 | 104.060000 | Wheat | 2025/3 | 10 | qPCR, 16S | Field |
| Qingdao | 36.130000 | 12.500000 | Wheat | 2023/5 | 10 | 16S | Lab |
| Shunyi_1 | 40.240000 | 116.710000 | Wheat | 2023/5 | 10 | 16S | Lab |
| Shunyi_2 | 40.240000 | 116.710000 | Wheat | 2024/4 | 10 | qPCR, 16S | Lab |
| Wenxian | 32.845268 | 104.877451 | Wheat | 2024/4 | 10 | 16S | Field |
| Wuhan | 30.500000 | 114.040000 | Barley | 2025/5 | 10 | PCR, 16S | Field |
| Xiangyang_1 | 32.011694 | 112.115833 | Wheat | 2024/4 | 10 | qPCR, 16S | Lab |
| Xiangyang_2 | 32.011694 | 112.115833 | Barley | 2025/5 | 10 | PCR, 16S | Field |
| Xianyang | 34.270000 | 108.700000 | Wheat | 2025/4 | 10 | PCR, 16S | Field |
| Xingtai | 37.061639 | 114.499750 | Wheat | 2024/5 | 10 | qPCR, 16S | Lab |
| Xiongan | 38.960000 | 116.040000 | Wheat | 2025/5 | 10 | PCR, 16S | Field |
| Yancheng | 32.980000 | 120.590000 | Barley | 2025/5 | 10 | PCR, 16S | Field |
| Yangling | 34.310995 | 108.105219 | Wheat | 2024/4 | 10 | 16S | Field |
| Yuncheng_1 | 35.027639 | 111.002139 | Wheat | 2023/5 | 10 | 16S | Lab |
| Yuncheng_2 | 35.027639 | 111.002139 | Wheat | 2024/5 | 10 | qPCR, 16S | Lab |
| Yuncheng_3 | 35.027639 | 111.002139 | Wheat | 2025/5 | 10 | PCR, 16S | Field |
| Zaozhuang_1 | 34.796694 | 117.257694 | Wheat | 2024/4 | 10 | qPCR, 16S | Lab |
| Zaozhuang_2 | 34.810000 | 117.320000 | Wheat | 2025/5 | 10 | PCR, 16S | Field |
| Zhengzhou | 34.766667 | 113.650000 | Corn | 2025/10 | 10 | qPCR, 16S | Field |
| Zhumadian_1 | 33.011583 | 114.016444 | Barley | 2024/5 | 10 | qPCR, 16S | Lab |
| Zhumadian_2 | 33.011583 | 114.016444 | Barley | 2025/4 | 10 | PCR, 16S | Field |
| Zhuozhou | 39.483028 | 115.968861 | Barley | 2024/5 | 10 | qPCR, 16S | Lab |

**Table 4.** Information for samples collected from Jutland (Denmark) as mapped in Figure 3. One sample (Aarhus_GD) was polymorphic for *Hamiltonella*, the other samples were all uninfected.

| *R. padi* sample | Latitude (degrees) | Longitude (degrees) | Field host plant | Date collected (year, month) | Aphids tested | Method | Number of lab generations before testing |
| --- | --- | --- | --- | --- | --- | --- | --- |
| Tilst | 56.19 | 10.12 | Barley | 2023/6 | 42 | qPCR, 16S | 6 |
| Orrild | 54.53 | 9.88 | Oats* | 2023/6 | 42 | qPCR, 16S | 5 |
| Aarhus_KT | 56.20 | 10.13 | Barley | 2023/7 | 42 | qPCR, 16S | 4 |
| Aarhus_GD | 56.20 | 10.11 | Barley | 2023/7 | 42 | PCR, qPCR, 16S | 4 |

\*Organic crop

**Table 5.** Information for *Sitobion avenae* collected from China as mapped in Figure 4 and a single sample of *Sitobion fragariae* collected from Australia.

| *Sitobion*<br>sample | Latitude<br>(degrees) | Longitude<br>(degrees) | Host Plant | Date<br>collected<br>(year,<br>month) | Aphids<br>tested | Method | Field or lab<br>reared?* |
| --- | --- | --- | --- | --- | --- | --- | --- |
| Changfeng | 32.27 | 117.09 | Barley | 2025/4 | 10 | PCR, 16S | Field |
| Changji | 44.36 | 86.5 | Wheat | 2025/4 | 10 | PCR, 16S | Lab |
| Dali | 25.67 | 100.3 | Barley | 2025/5 | 10 | PCR, 16S | Field |
| Ezhou | 30.01 | 114.91 | Wheat | 2025/4 | 10 | PCR, 16S | Field |
| Gushi | 36.03 | 106.24 | Wheat | 2025/5 | 10 | 16S | Field |
| Guyuan | 36.03 | 106.24 | Wheat | 2025/5 | 10 | 16S | Field |
| Haidian_1 | 39.95 | 116.29 | Wheat | 2025/4 | 10 | qPCR, 16S | Field |
| Haidian_2 | 39.95 | 116.29 | Wheat | 2025/9 | 10 | PCR, qPCR,<br>16S | Lab |
| Handan | 36.55 | 114.41 | Wheat | 2025/5 | 10 | qPCR, 16S | Field |
| Hangzhou | 30.24 | 120.2 | Wheat | 2025/4 | 10 | qPCR, 16S | Field |
| Hefei | 31.82 | 117.22 | Barley | 2025/4 | 10 | PCR, 16S | Field |
| Jiangyou_1 | 31.89274 | 104.78822<br>7 | Wheat | 2024/4 | 10 | PCR, 16S | Field |
| Jiangyou_2 | 31.89270<br>1 | 104.78825<br>2 | Wheat | 2024/4 | 10 | PCR, 16S | Field |
| Jiangyou_3 | 31.89573<br>4 | 104.78775<br>6 | Wheat | 2024/4 | 10 | PCR, 16S | Field |
| Jiangyou_4 | 31.89040<br>5 | 104.78931<br>2 | Wheat | 2024/4 | 10 | PCR, 16S | Field |
| Jiangyou_5 | 31.89387<br>7 | 104.78834 | Wheat | 2024/4 | 10 | PCR, 16S | Field |
| Jiangyou_6 | 31.89657 | 104.78697<br>9 | Wheat | 2024/4 | 10 | PCR, 16S | Field |
| Jiangyou_7 | 31.89676<br>9 | 104.78745<br>5 | Wheat | 2024/4 | 10 | PCR, 16S | Field |
| Jiangyou_8 | 31.89255<br>8 | 104.78808<br>5 | Wheat | 2024/4 | 10 | PCR, 16S | Field |
| Jiangyou_9 | 31.89612 | 104.78757<br>6 | Wheat | 2024/4 | 10 | PCR, 16S | Field |
| Jiangyou_10 | 31.89657<br>2 | 104.78770<br>1 | Wheat | 2024/4 | 10 | PCR, 16S | Field |
| Wuhan | 30.5 | 114.04 | Barley | 2025/4 | 10 | PCR, 16S | Field |
| Xiangyang | 32 | 112.12 | Barley | 2025/4 | 10 | PCR, 16S | Field |
| Xianyang | 34.27 | 108.7 | Wheat | 2025/5 | 10 | PCR, 16S | Field |
| Xingtai | 37.07 | 114.5 | Wheat | 2025/5 | 10 | PCR, 16S | Field |
| xiongan | 38.96 | 116.04 | Wheat | 2025/5 | 10 | PCR, 16S | Field |
| Yuncheng | 35.3 | 111.00 | Wheat | 2025/5 | 10 | qPCR, 16S | Field |
| Zaozhuang | 34.81 | 117.32 | Wheat | 2025/5 | 10 | qPCR, 16S | Field |
| Bundoora* | -37.724 | 145.0546 | Grass | 2025/12 | 10 | qPCR, 16S | Field |

\**S. fragariae* collection from Australia

**Figure 1.**
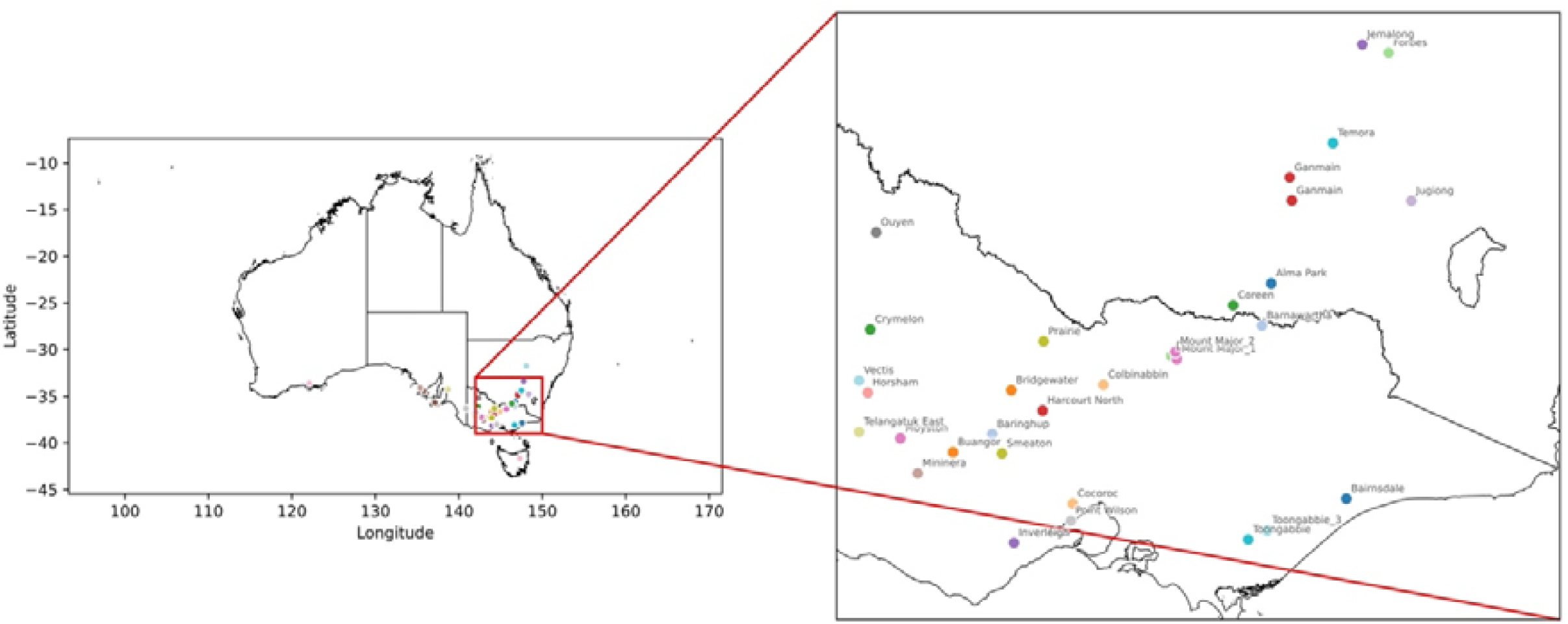
Distribution of Australian samples with most coming from Victoria and New South Wales (inset). Details of locations are provided in Table 2.

**Figure 2.**
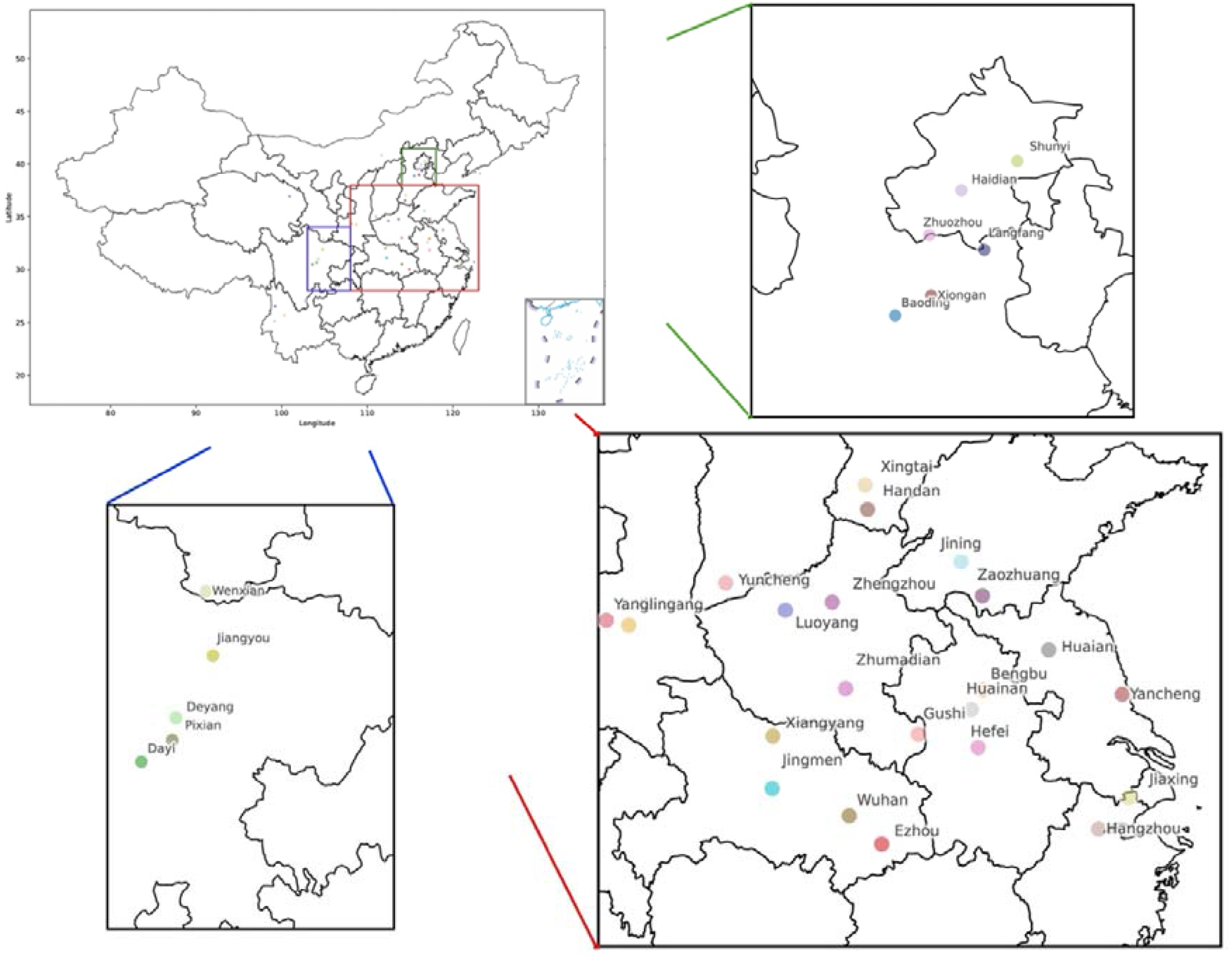
Distribution of Chinese *R. padi* samples (map and insets). Details of locations are provided in Table 3.

**Figure 3.**
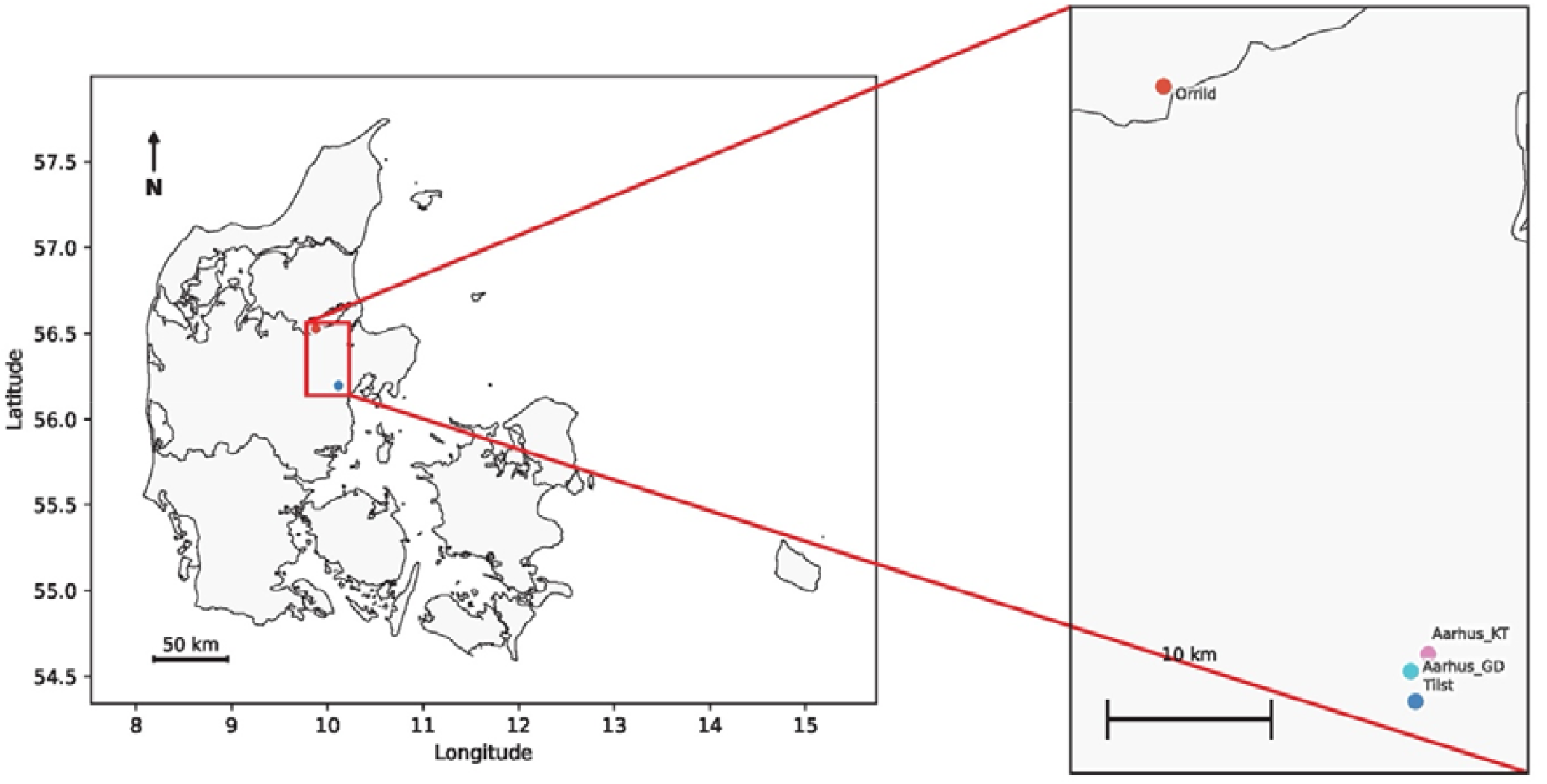
Distribution of *R. padi* samples listed in Table 4 from Jutland (Denmark).

### 16S rRNA gene metabarcoding and quantitative PCR

We used DNA metabarcoding to characterise the microbiome of all *R. padi* and *Sitobion* sp. samples. Individuals were pooled to provide sufficient DNA for next generation sequencing. Two replicate DNA extractions, each containing a pool of 5 individuals, were performed for each sample using a DNeasy® Blood & Tissue kit (Qiagen, Hilden, Germany). Metabarcoding targeted the hypervariable V3-V4 region of the bacterial 16S rRNA gene and was carried out by Novogene (Novogene, HK, Co. Ltd), using the universal primers 341F and 806R. Sequence analysis was performed using a standard QIIME2 pipeline [31], according to methods described previously (Yang et al., 2023a).LFN (low-frequency noise) filters were used to discard variants with low read counts and sequencing contamination (Yang et al., 2023a). An average of 212,605 reads per sample were retained after each of the quality filtering and assembly steps from our 16S metabarcoding data.

Quantitative PCR (qPCR) assays were used to screen for secondary endosymbionts in *R. padi* to validate 16S rRNA gene metabarcoding results or when rapid screening was required for culture maintenance. qPCR assays were performed using a LightCycler® 480 High Resolution Melting Master kit (Roche, New South Wales, Australia) and IMMOLASE™ DNA polymerase (5 U/µl, Bioline). Amplification conditions consisted of an initial 10 min pre-incubation at 95°C (ramp rate = 4.8°C/s), followed by 40 cycles of 95°C for 5 s (4.8°C/s), 58°C for 15 s (2.5°C/s), and 72°C for 30 s (4.8°C/s). Two primer sets were used to amplify markers targeting aphid β*-actin* as a reference gene and the target endosymbiont (Table 6). Crossing point (Cp) values from two or three consistent technical replicates were averaged. Relative endosymbiont densities were estimated using the 2^-Δ*Cp*^ method, where Δ*Cp* represents the difference between the Cp values of the target endosymbiont and the β*-actin* reference gene.

**Table 6.**
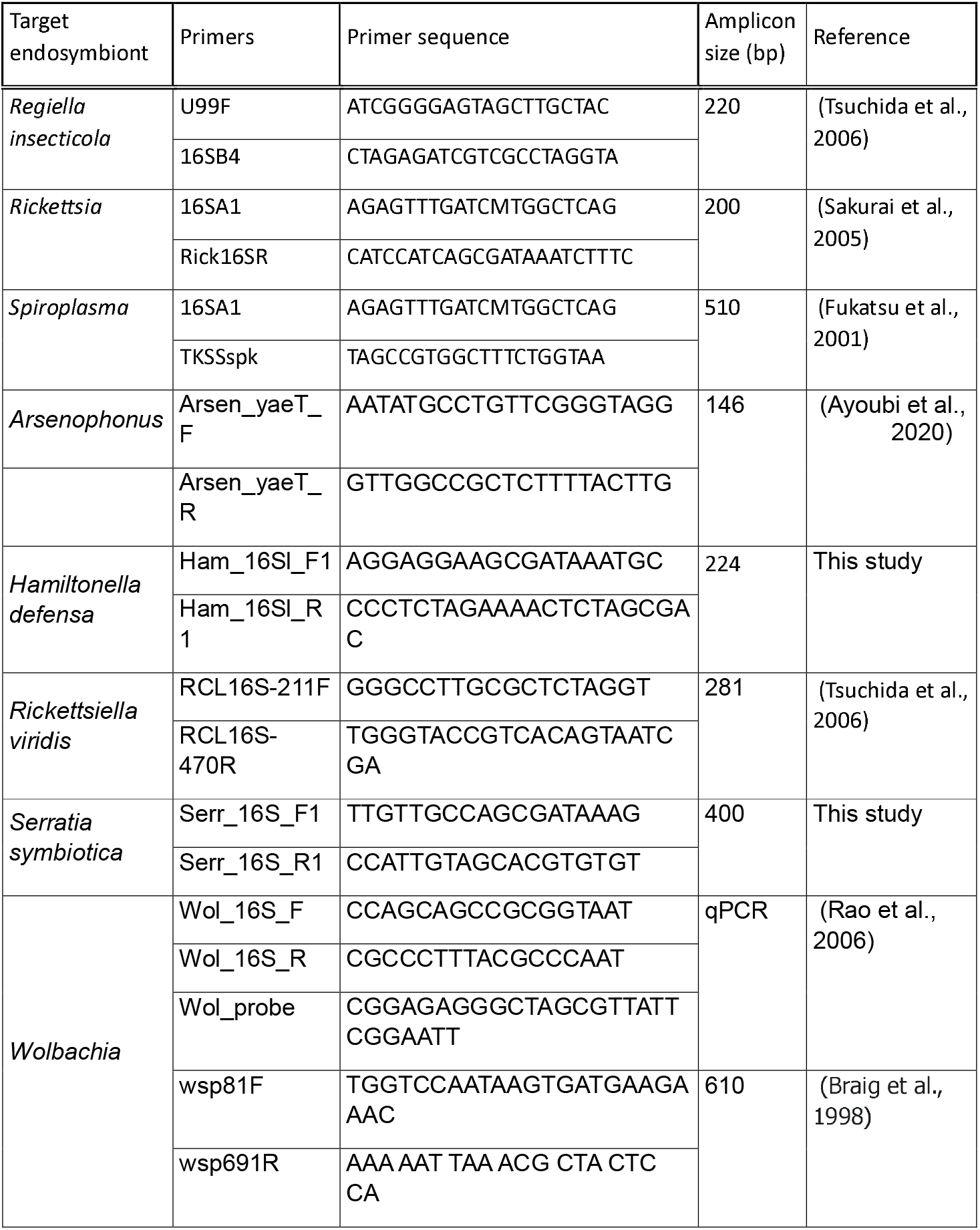
Primers for endosymbiont detection using conventional PCR or qPCR.

### Conventional PCR and Sanger sequencing

Presence of the endosymbiont amplicon was confirmed by running a standard PCR using the same program and primers as used for qPCR. We prepared a final volume of 25 µl for each PCR reaction mixture. Each reaction contained 14.85 µl of PCR-graded water (Elga Purelab® flex), 1.25 µl of each forward primer and reverse primer (10μM), 1.25 µl MgCl2 (50 mM, Bioline), 2.5 µl of 10× Standard ThermoPol® Reaction Buffer (B9015S, NEB), 2.0 µl dNTPs (2.5 mM), 0.4 µl of IMMOLASE™ DNA polymerase (5 U/µl, Bioline), and 1.5 µl of DNA template. We run 10 µl of PCR product on a 2% molecular biology grade agarose gel (Scientifix, Victoria, Australia) and observing a clear band with appropriate size for each primer pair. Remaining PCR products were then sent for Sanger sequencing (Macrogen, Inc., Seoul, South Korea). Sequencing chromatograms were examined and processed with Geneious 9.18 software.

## Results

### Literature review

We identified 9 papers with information on secondary endosymbionts in *R. padi* covering 10 sampling collections (Table 1). In some cases only pooled samples were tested from different locations whereas in other cases individuals were sampled so that frequencies could be computed. A diversity of endosymbionts was detected in two studies (Guo et al., 2019; Majeed et al., 2022) including one where two regions were surveyed. In contrast, other studies failed to detect endosymbionts or detected them at a very low incidence (<0.03% in the case of Li et al (2025) despite the ability to screen for a lot of bacterial types and wide geographic areas covered (Donner et al., 2024; Fakhour et al., 2018; Yang et al., 2023a). In a few cases, only one secondary endosymbiont was detected, with *Hamiltonella* being detected at a substantial frequency in one study (Leybourne et al., 2020). Studies varied in the nature of the controls applied and whether multiple approaches were adopted in validating the presence of endosymbionts (Table 1).

### New Australian samples

We failed to detect any secondary endosymbionts in the 48 Australian samples sourced from grasses or cereals and tested by 16S (Figure 5) and/or qPCR. *Buchnera aphidicola* was detected in all samples tested with 16S and validated by PCR or qPCR. We also collected one sample of *Sitobion fragariae*. This sample was infected by *Regiella* as determined by both 16S and qPCR screening.

**Figure 4.**
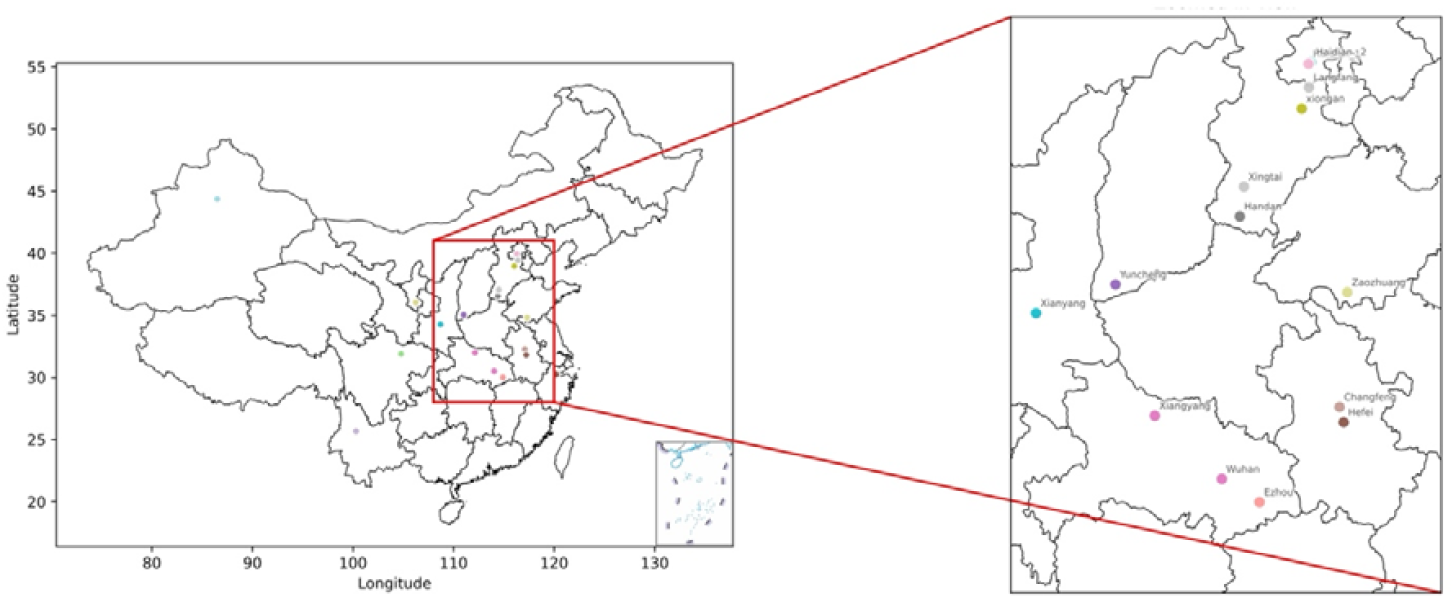
Distribution of *Sitobion avenae* samples collected from China.

**Figure 5.**
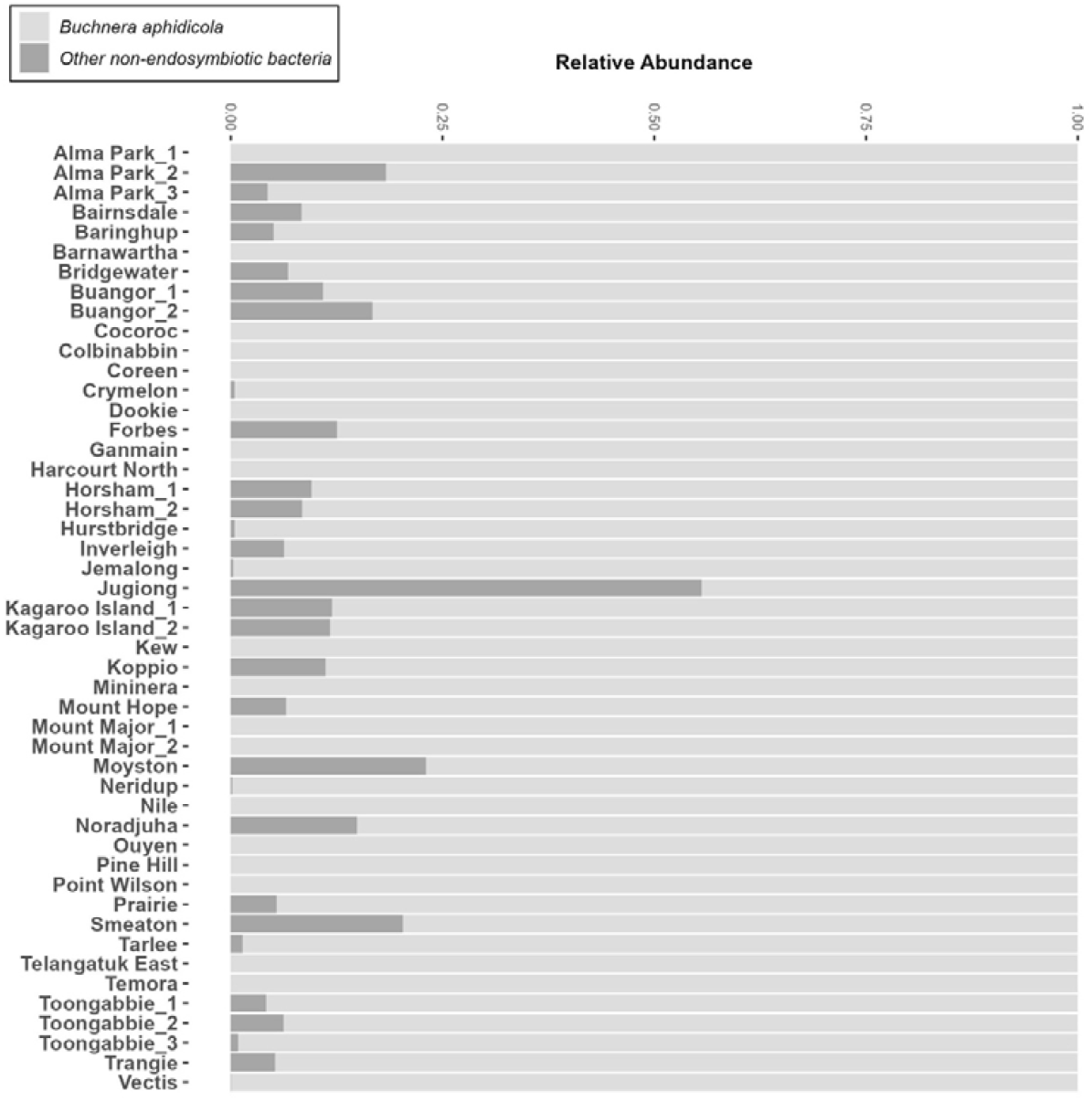
Relative abundance of endosymbionts detected in Australian *R. padi* samples with 16S and validated by PCR or qPCR. The x-axis represents the mean relative abundance of Illumina reads per sample, while the y-axis lists each sample. Bacterial taxa are indicated in the legend.

### New Chinese samples

We failed to detect any secondary endosymbionts in the 59 Chinese samples tested by 16S rRNA gene sequencing (Figure 6) and qPCR, irrespective of whether samples came from wheat, barley or corn, and whether they came directly from the field or tested after a period of lab culture. Note that this includes several samples that were repeatedly collected across several years from the same location (Table 3). As for the Australian samples, *Buchnera aphidicola* was detected in all samples tested with 16S.

**Figure 6.**
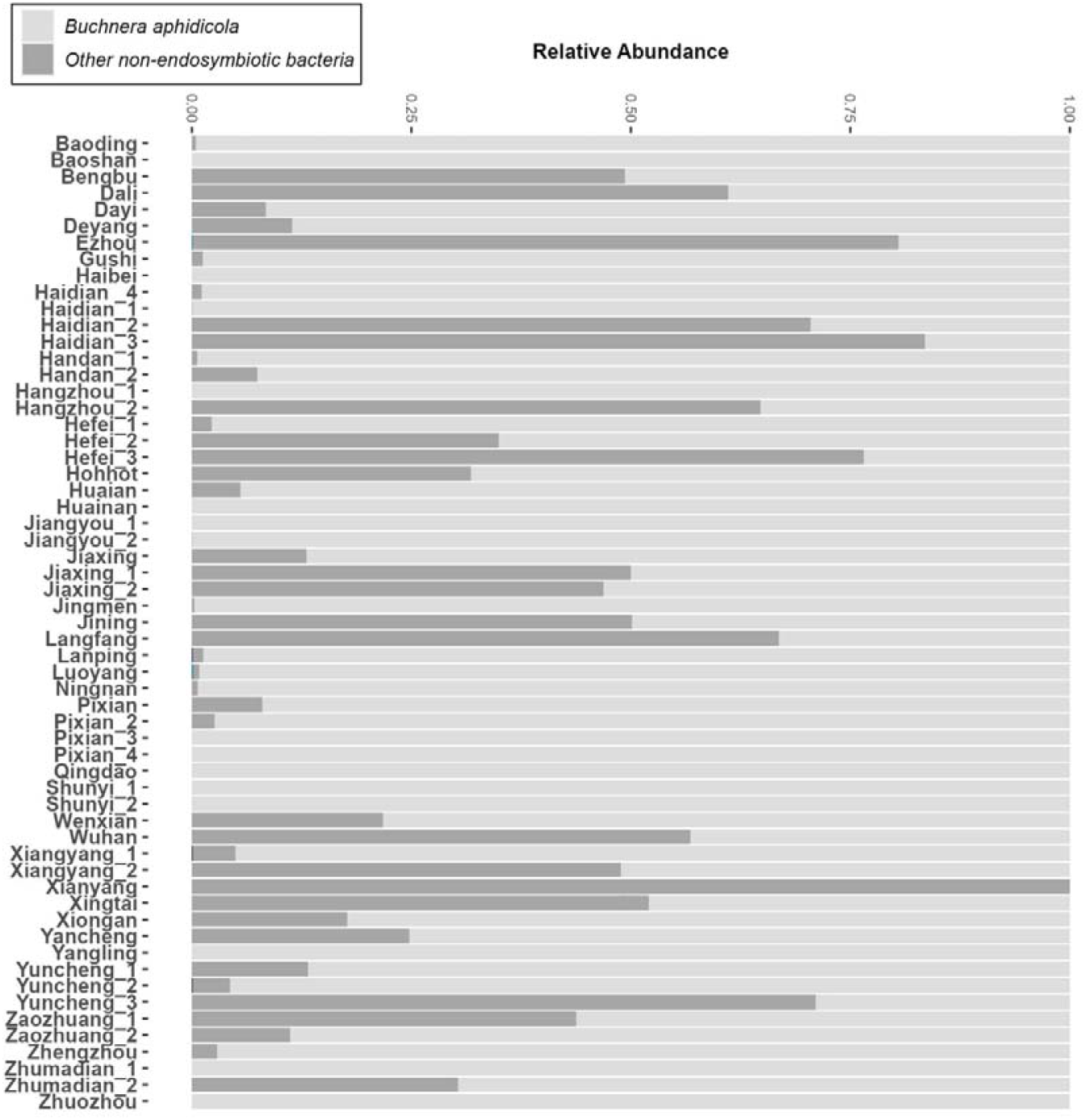
Relative abundance of endosymbionts in Chinese *R. padi* samples with 16S rRNA gene sequencing and validated by PCR or qPCR. Bacterial taxa are indicated in the legend.

We also tested 28 samples of *Sitobion avenae* collected at the same time as the *R. padi* samples. These proved to be commonly infected with *Regiella insecticola* (Figure 7). We also recorded one sample infected with *Spiroplasma* and three samples infected with *Sitobion miscanthi* L-type symbiont (SMLS) which has been linked to the endosymbiont genus Candidatus *Hemipteriphilus* (Li et al., 2016; Li et al., 2023). We note that *Buchnera aphidicola* was not found in ten samples of *S. avenae* which requires further investigation.

**Figure 7.**
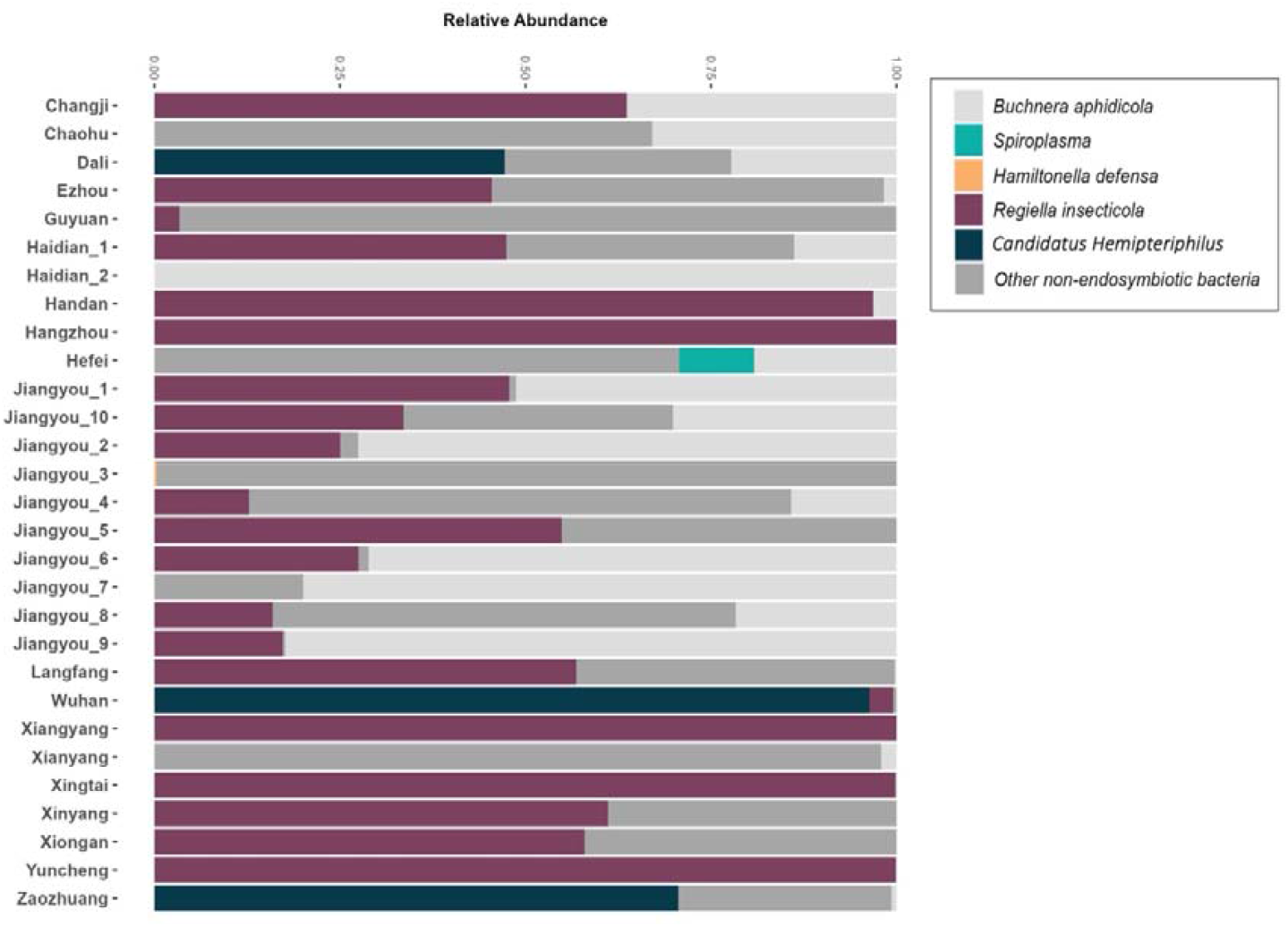
Relative abundance of endosymbionts in Chinese *S. avenae* samples with 16S rRNA gene sequencing and validated by PCR or qPCR. Bacterial taxa are indicated in the legend.

### Danish samples

Aphids were sampled from four cereal crops after different periods in the laboratory (Table 4). Three of these were uninfected (Figure 8) but *Hamiltonella defensa* was detected in around half of the aphids from one sample when aphids were tested by qPCR. Subsequently, a line completely infected by this endosymbiont was generated and it has remained stably infected for 40 generations. The infection was confirmed by PCR.

**Figure 8.**
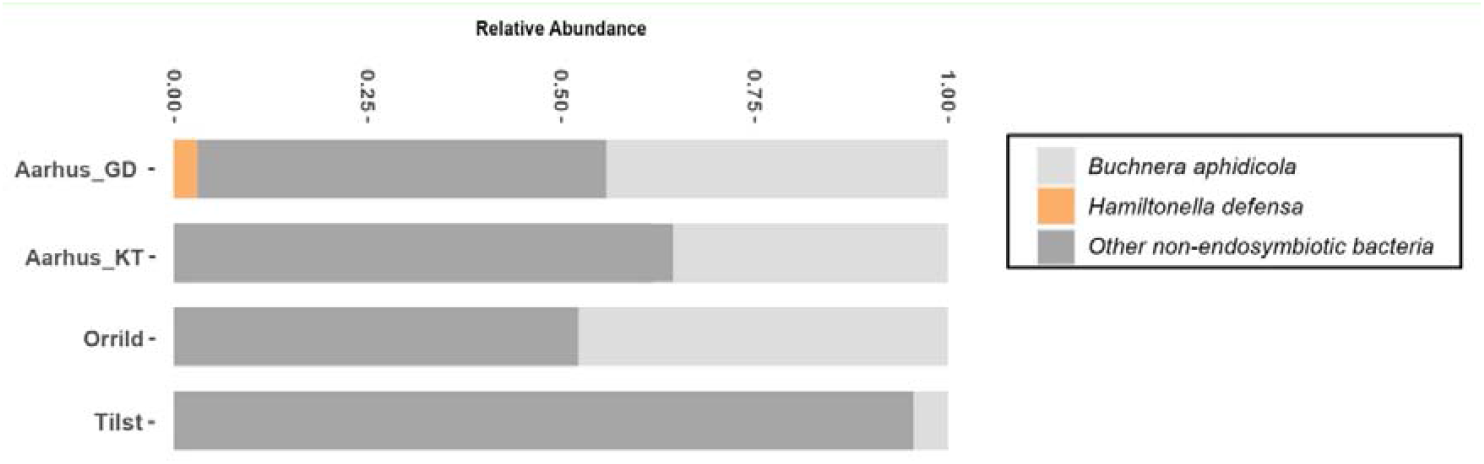
Relative abundance of endosymbionts in Danish *R. padi* samples with 16S rRNA gene sequencing and validated by PCR or qPCR. Bacterial taxa are indicated in the legend.

#### Evidence for false-positive detection due to inadequate validation and contamination

To validate the endosymbiont detections from 16S rRNA gene sequencing, all aphid samples were screened before or after sequencing using conventional PCR and/or qPCR assays targeting specific symbionts. Detections were considered conclusive only when confirmed across the methods used. In most cases, the results were consistent among the methods. However, false-positive detections were occasionally observed with conventional PCR amplification. In several cases, PCR amplification produced bands of the expected size on agarose gels (Figure 9A-C), which would typically be interpreted as positive detections. However, Sanger sequencing of these amplicons showed that they did not correspond to the expected endosymbionts (Supplementary material). Specifically, sequences obtained from putative *Regiella* amplicons matched *Acinetobacter lwoffii*, while products amplified with *Spiroplasma* primers matched *Planococcus sp*. Similarly, amplicons generated using *Rickettsia*-specific primers were identified as *Agrobacterium fabrum*. These results demonstrate that PCR amplification alone, even when producing bands of the expected size, can lead to incorrect identification of endosymbionts if not validated by DNA sequencing.

**Figure 9.**
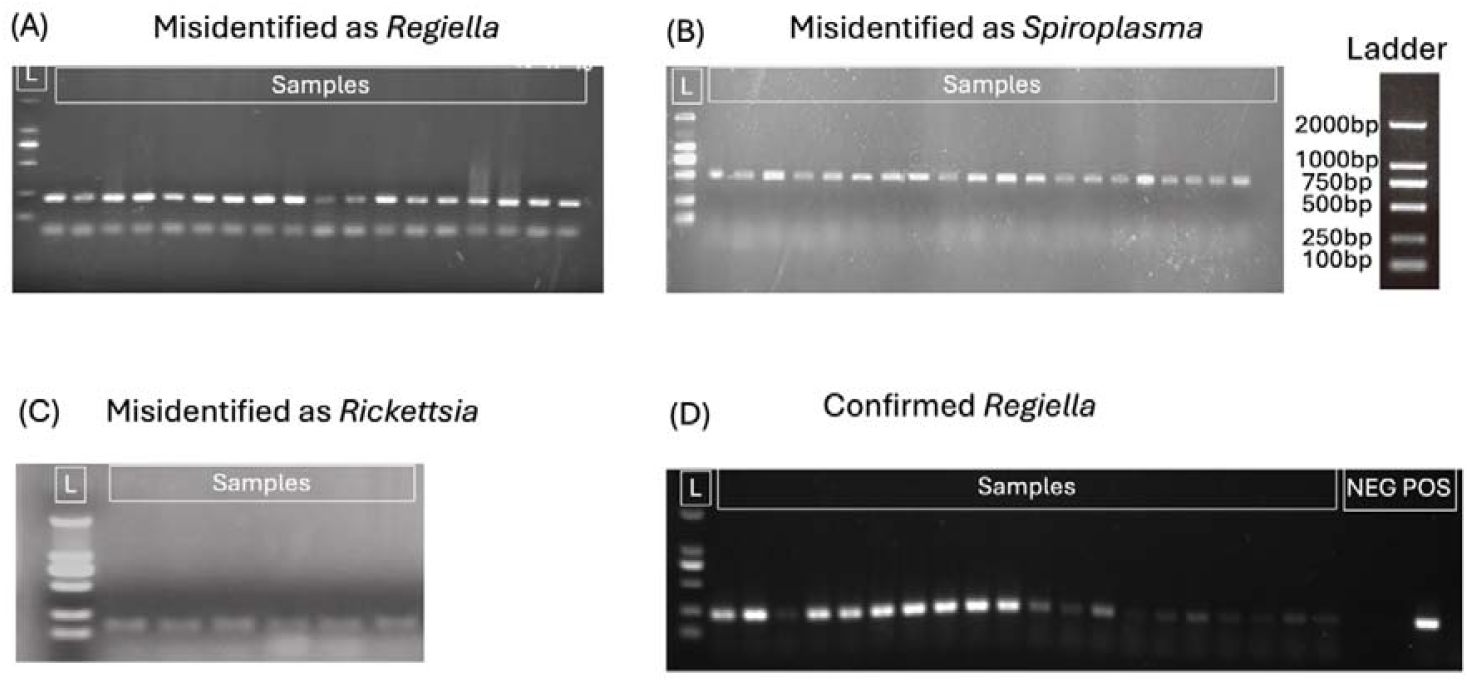
Positive PCR amplification does not necessarily confirm endosymbiont presence in *R. padi*. (A-C) Conventional PCR screening for three endosymbionts in individual *R. padi* samples collected from Pixian, China (2024), showing the current band size for *Regiella* (A), *Spiroplasma* (B) and *Rickettsia* (C). (D) Conventional PCR screening for *Regiella* in *R. padi* samples collected from Baoding, China (2025), including negative (NEG) and positive (POS) controls, with *Regiella* confirmed in several samples but contamination present.

Further evidence for unreliable detection was obtained through additional screening assays targeting *Regiella* (Figure 9D). Negative and positive controls were included in these conventional PCR assays, indicating that the reactions themselves were expected to have been successful. Nevertheless, when all samples that initially produced bands were re-screened using independent assays (including 16S rRNA gene sequencing – see Figure 6), none were confirmed as being infected with *Regiella* (Figure S1) suggesting contamination. These results suggest that erroneous detections can occur even when appropriate controls are included.

Together, these findings highlight important methodological limitations associated with PCR-based detection of endosymbionts. In particular, reliance on expected amplicon size without sequence validation, as well as the presence of contamination, can lead to false-positive detections. These issues underscore the importance of sequence confirmation and robust validation steps when assessing endosymbiont presence in aphid populations.

## Discussion

Clearly our results revealing little evidence for presence of secondary endosymbionts stand in contrast to the high diversity of endosymbionts detected in *R. padi* in two previous studies from China (Guo et al., 2019; Majeed et al., 2022) and one from Europe (Guo et al., 2019). Our surveys in both China and Australia were aimed at being comprehensive and so involved a large number of samples that were screened using multiple validation approaches. Together with the additional sequencing and re-screening assays conducted here, our results suggest that facultative endosymbionts are genuinely rare in *R. padi* populations. This raises the question whether there has been a dramatic change in endosymbiont diversity across time/crops or that some previous detections were erroneous.

We note that *Hamiltonella* was detected in a sample from Denmark and was also detected in the UK (Leybourne et al., 2020) and in the Netherlands (Donner et al., 2024) so we suspect that this is a reliable result, and that this symbiont is likely a genuine, though apparently uncommonly associated with *R. padi*. However, the detection of a high frequency of other endosymbionts in the other studies (Table 1) remains a mystery particularly as some positive and negative controls were included in the Guo et al. (2019) study. However, we note that only one detection approach was used in that study.

We do not believe that host plant species explains the previously reported high endosymbiont diversity in *R. padi*, as we deliberately sampled maize/corn *R. padi* which was the source of one of the high diversity reports (Guo et al., 2019). Instead, contamination from other aphid species may provide a more likely explanation. In particular, *Sitobion avenae*, which was frequently collected alongside *R. padi*, carried multiple endosymbionts, including *Regiella*. This is consistent with previous surveys reporting *Regiella* and other facultative symbionts in *S. avenae* (Fakhour et al., 2018; Hu et al., 2020; Łukasik et al., 2013). Importantly, when samples that initially produced positive PCR bands were re-screened using independent assays under stricter experimental conditions, none were confirmed to be infected with *Regiella* (Figure S1), suggesting that contamination can be effectively eliminated through robust validation procedures and that convincing absence results can be obtained. We are unaware of previous surveys that include *S. fragariae*, although this was the only *Sitobion* species that we were able to collect locally in Australia.

The other bacteria identified in these false-positive amplicons are also commonly associated with insects or plants. For example, *Acinetobacter lwoffii*, amplified using putative *Regiella* primers, is commonly reported from aphids (Yandigeri et al., 2024) and other insects (Karut et al., 2020). Similarly, amplicons generated using *Rickettsia*-specific primers matched *Agrobacterium fabrum*, a widespread plant-associated bacterium, while products amplified using *Spiroplasma*-specific primers matched *Planococcus* sp., a bacterial genus associated with hydrocarbon degradation (Waghmode et al., 2020). PCR-based approaches were generally reliable, with most detections being consistent across conventional PCR, qPCR, and 16S rRNA gene sequencing. However, our results highlight the importance of sequence confirmation and independent validation, particularly when screening for low-abundance or previously unreported endosymbionts.

The apparent rarity of secondary endosymbionts in *R. padi* may be biologically informative. Unlike several other cereal aphids, such as *Sitobion avenae*, which commonly harbour multiple facultative symbionts, *R. padi* may possess ecological, physiological, or evolutionary characteristics that limit the establishment or persistence of these bacteria. In this sense, it matches other important pest aphids that also mostly lack secondary endosymbionts including the green peach aphid (Yang et al., 2023b) and Russian wheat aphid (Swanevelder et al., 2010). Another message from our study is that when carrying out surveys, it is important to validate the results in several ways which should particularly include additional sequencing of the putative endosymbionts. In this context it could be useful to isolate lines fixed for the endosymbiont in the laboratory so that additional tests can be done, although it is also possible that endosymbionts are lost during laboratory culture (Yang et al., 2023a).

## Supporting information

BLASTn analysis

## Acknowledgments

This work was undertaken as part of the Australian Grains and Horticulture Pest Innovation Program (AGHPIP), supported through funding provided by the Grains Research and Development Corporation (UOM1906-002RTX; UOM2404-006RT), and the Australian Horticultural Pest Innovation Program (ST23002) funded through Hort Innovation Frontiers with co-investment from The University of Melbourne and Cesar Australia and contributions from the Australian Government. The survey in Denmark was supported by Villum Fonden (40841), and the survey in China was supported by the China Agriculture Research System (CARS-05-03A-16) and the Scientific and Technological Achievement Transformation Project of the Sichuan Academy of Agricultural Sciences (2025ZSSFGH12; 2026ZSSFGH21). We thank the numerous people who assisted with aphid collections, including Evatt Chirgwin, Kaya Moore, Rob Harrod, Lilia Jenkins, Leo McGrane, Elia Pirtle, Piotr Trebicki, Aston Arthur, Nigel Myers, Adriana Arturi, Brad Mills, Ben Lenehan, Jim Cronin, Rodney Krueger, Nigel Myers, Stephanie Veskoukis, Kara Bryant, Sam Ward and Dave Cox. We are also grateful to Jade Russell and Mel Berran for assistance with aphid culturing and preservation, and to Monica Stelmach, Kelly Richardson, Mason Mason, Yifei Jin and Yuheng Chen for technical assistance.

