## Supplementary Figure 1 for "Revisiting the diversity of secondary endosymbionts in the major pest oat aphid, *Rhopalosiphum padi*"

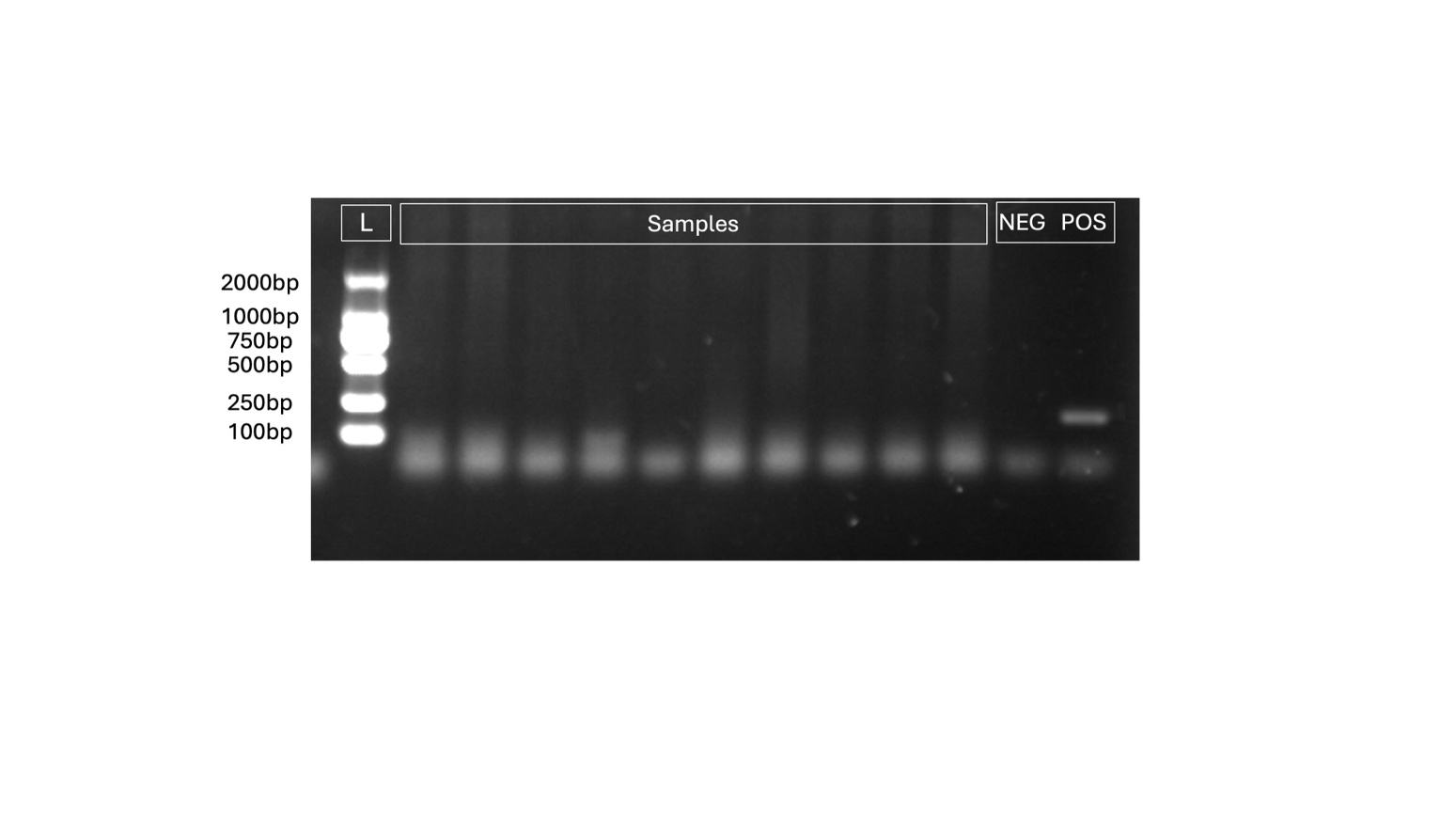

Figure S1. Re-screening assays for putative *Regiella*-positive R. padi samples. Conventional PCR assays targeting Regiella were repeated on samples that initially produced bands of the expected size. Negative (NEG) and positive (POS) controls were included to verify PCR performance. None of the re-screened samples produced convincing amplification consistent with Regiella infection, suggesting that the initial detections were false positives, likely resulting from contamination or non-specific amplification.
